# Characterization of cognitive-motor function in women who have experienced intimate partner violence-related brain injury

**DOI:** 10.1101/2021.01.29.428699

**Authors:** Naomi Maldonado-Rodriguez, Clara Val Crocker, Edward Taylor, K. Elisabeth Jones, Krystal Rothlander, Jon Smirl, Colin Wallace, Paul van Donkelaar, on behalf of the Canadian Traumatic Brain Injury Research Consortium

**Author notes:** Corresponding Author: Paul van Donkelaar, School of Health and Exercise Science, 1147 Research Rd., University of British Columbia Okanagan, Kelowna, BC, V1V 1V7. Naomi Maldonado-Rodriguez and Clara Val Crocker contributed equally to this work and share first authorship on this manuscript.

## Abstract

Intimate partner violence (IPV) affects at least 1 in 3 women worldwide and up to 92% report symptoms consistent with brain injury (BI). Although a handful of studies have examined different aspects of brain structure and function in this population, none has characterized potential deficits in cognitive-motor function. This knowledge gap was addressed in the current study by having participants who had experienced IPV complete the bimanual Object Hit & Avoid (OHA) task on a Kinarm End-Point Lab. BI load, as assessed by the Brain Injury Severity Assessment (BISA) tool as well as measures of comorbidities (PTSD, anxiety, depression, substance use, history of abuse) were also collected. Results demonstrated BI load accounted for a significant amount of variability in the number of targets hit and average hand speed. PTSD, anxiety, and depression also contributed significantly to the variability in these measures as well as to the number and proportion of distractor hits, and the object processing rate. Taken together, these findings suggest IPV-related BI, as well as comorbid PTSD, anxiety, and depression, disrupt the processing required to quickly and accurately hit targets while avoiding distractors. This pattern of results reflects the complex interaction between the physical injuries induced by the episodes of IPV and the resulting impacts these experiences have on mental health.

## INTRODUCTION

It is estimated one in three women will experience intimate partner violence (IPV) in their lifetime.^1^ IPV is considered one of the most prevalent forms of violence against women and involves physical, sexual, and/or emotional abuse, as well as controlling behaviours from an intimate partner.^1^ It leads to a multitude of physical, psychological, neurological, cognitive, and emotional consequences, most notably, anxiety, depression, and post-traumatic stress disorder (PTSD).^2–5^ However, it is increasingly recognized brain injury (BI) resulting from violent blows to the head, face, and neck, and/or strangulation, is also a common part of this experience.^6,7^ Even though some survivors may experience neuropsychological impairment without BI^8^, a recent review^9^ reported up to 92% of women reported symptoms consistent with BI following an IPV incident – including headaches, memory loss, difficulty thinking, paying attention, or getting organized.^4,10^ Thus, it is clear that IPV may result in a multitude of physical, psychological, and social consequences, however, the extent to which BI plays a role in such dysfunction is unknown.

There is a small body of research directly examining the relationship between IPV-related BI and brain structure and function. In particular, Valera and colleagues have shown BI severity is related to decreases in i) memory, learning, and cognitive flexibility^11^; ii) functional connectivity in the default mode network similar to that observed in individuals with BI from other causes^3^; and iii) functional anisotropy indicating white matter disruptions in the posterior and superior corona radiata^5^. Importantly, these relationships were present even when taking psychopathological comorbidities (e.g., PTSD, anxiety, depression) into account. These studies have provided much-needed evidence demonstrating the symptoms reported by women experiencing IPV-related BI are more than simply the result of mental health disorders and have provided quantitative evidence of BI in this population.

BIs and repetitive subconcussive head impacts in other populations (e.g., contact sport athletes, military personnel) are often associated with alterations in various aspects of neurocognitive function.^12–17^ Given the chronic and repetitive nature of IPV, it is likely that alterations in neurocognitive function may also be present in survivors of IPV-related BI. Toward this end, the purpose of the current study was to characterize a specific aspect of neurocognition – cognitive-motor function – in a cohort of women who have experienced IPV and examine the extent to which it was related to clinical measures of executive function. We hypothesized that IPV survivors who reported more exposure to head impacts and strangulation during episodes of IPV would display more pronounced cognitive-motor deficits relative to those with less exposure.

## METHODS

### Participants

Women who had experienced IPV were recruited from a local women’s shelter and other women-serving organizations. Individuals were eligible if they identified as women, were between 18-50 years of age, and had previously experienced IPV. Exclusion criteria included being pregnant, having experienced non-partner related BI in the last 6 months, having been diagnosed with a neurological or neuropsychiatric disorder other than BI that could affect cerebrovascular, neurocognitive, and/or sensorimotor function (e.g. stroke, Parkinson’s, Alzheimer’s, migraine, seizures, schizophrenia) or an orthopaedic, musculoskeletal, or degenerative disorder that could affect balance control (e.g. osteoarthritis, recent lower limb joint surgery). Participants provided written informed consent and the study was approved by the University of British Columbia Clinical Research Ethics Board.

### Procedures

This study is part of a large community based-research project focused on IPV-related BI (*SOAR: Supporting Survivors of Abuse and Brain Injury through Research)* in which participants completed two sessions separated by a minimum of 3 days. Participants were asked to refrain from consuming alcohol or using drugs for 24 hours prior to each session. During the first session, participant demographics (including age, years of education, ethnicity, indigeneity, access to housing, and time since last injury), CAPS-IV, Beck depression and anxiety inventories, women’s experiences with battering scale^17^, and initial substance use scale^18^ were collected. In addition, the Brain Injury Severity Assessment (BISA)^3,5,10,11^ was used to characterize BI load resulting from impacts targeting the head, face, and neck and strangulation that resulted in alterations of consciousness during episodes of IPV as well as from non-partner related BI events (e.g., falls, car accidents, etc). The BISA yields an overall score of 0-8 comprised of recency, frequency, and severity subcomponents.

During the second visit, participants completed a series of physiological, neurocognitive, and sensorimotor assessments. This study reports the results for the Object Hit and Avoid (OHA) task performed on the Kinesiological Instrument for Normal and Altered Reaching Movement (KINARM) End-Point Lab (BKIN Technologies, Kingston, Ontario, Canada). The KINARM consists of two moveable robotic arms, one for each hand, and is equipped with a display screen that provides feedback during a variety of sensorimotor tasks. The reliability, sensitivity, and objectivity of the KINARM technology to detect motor, sensory, and cognitive deficits in brain injured participants have been previously described.^20–22^ In the OHA task, participants were asked to hit two predetermined target shapes while avoiding all other distractor shapes. Over the course of the 2-minute task, the shapes descended from the top of the screen with increasing speed, starting at ~10 cm/s and ending at ~50 cm/s. Performance on the task was assessed via the parameters outlined in Table 1.^23^ Finally, to determine if OHA task performance was related to clinical measures of executive function, participants also completed the Trail-Making A and B Tasks (TMT) and the Behavior Rating Inventory of Executive Function – Adult (BRIEF-A).

**Table 1.**
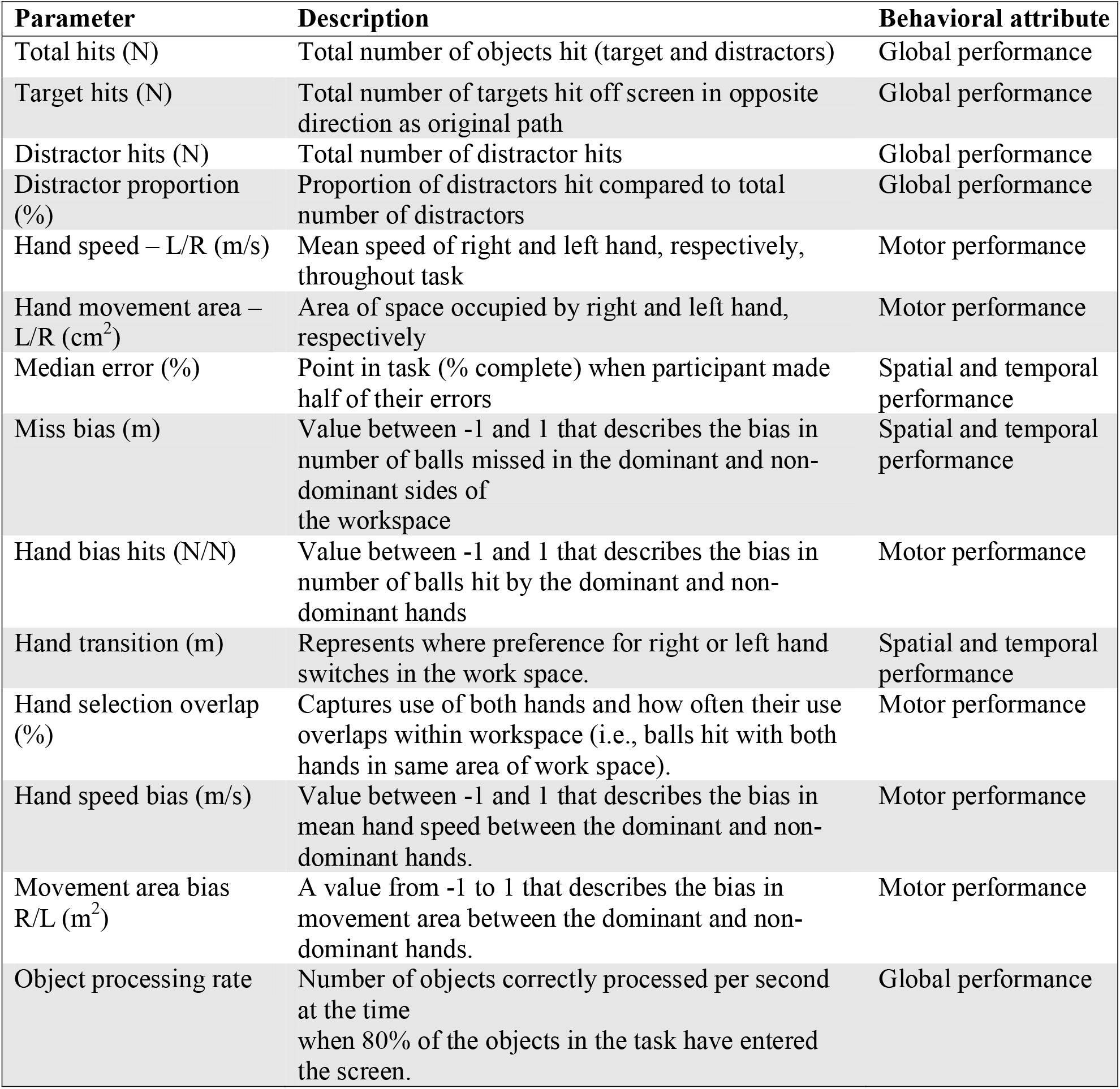
Task parameters for the Object Hit Avoid Task and associated behavioral attributes

| Parameter | Description | Behavioral attribute |
| --- | --- | --- |
| Total hits (N) | Total number of objects hit (target and distractors) | Global performance |
| Target hits (N) | Total number of targets hit off screen in opposite direction as original path | Global performance |
| Distractor hits (N) | Total number of distractor hits | Global performance |
| Distractor proportion (%) | Proportion of distractors hit compared to total number of distractors | Global performance |
| Hand speed – L/R (m/s) | Mean speed of right and left hand, respectively, throughout task | Motor performance |
| Hand movement area – L/R (cm<sup>2</sup>) | Area of space occupied by right and left hand, respectively | Motor performance |
| Median error (%) | Point in task (% complete) when participant made half of their errors | Spatial and temporal performance |
| Miss bias (m) | Value between -1 and 1 that describes the bias in number of balls missed in the dominant and non-dominant sides of the workspace | Spatial and temporal performance |
| Hand bias hits (N/N) | Value between -1 and 1 that describes the bias in number of balls hit by the dominant and non-dominant hands | Motor performance |
| Hand transition (m) | Represents where preference for right or left hand switches in the work space. | Spatial and temporal performance |
| Hand selection overlap (%) | Captures use of both hands and how often their use overlaps within workspace (i.e., balls hit with both hands in same area of work space). | Motor performance |
| Hand speed bias (m/s) | Value between -1 and 1 that describes the bias in mean hand speed between the dominant and non-dominant hands. | Motor performance |
| Movement area bias R/L (m<sup>2</sup>) | A value from -1 to 1 that describes the bias in movement area between the dominant and non-dominant hands. | Motor performance |
| Object processing rate | Number of objects correctly processed per second at the time when 80% of the objects in the task have entered the screen. | Global performance |

### Statistical Analysis

Stepwise multiple regression analyses were completed in R (Version 3.6.3) to evaluate the relationship between BI load and task performance while accounting for potential confounders (i.e., age, history of non-partner related BI, PTSD, depression, anxiety, history of abuse, substance use). Model fits were assessed using Akaike Information Criterion (AIC). Where appropriate, post-hoc analyses were completed using t-tests. Significance was set at *p* < .05. To better understand whether OHA performance measures predicted by the BISA scores were related to clinical measures of executive function (TMT, BRIEF-A), we calculated the correlations between these variables.

## RESULTS

### Demographics

Forty women participated in the study. Table 2 outlines the demographic characteristics of our sample and the scores on the assessments of potential confounders. As expected, the experience of a large majority of participants fulfilled the criteria for battering according to the WEB scale (defined as scores > 20). Many also exhibited elevated levels of depression (62%), anxiety (58%), and PTSD (95%). Moreover, a substantial proportion, (75%) had experienced at least 1 non-partner related BI. Finally, 22.5% of the participants were Indigenous women. This is an overrepresentation of the general population of the area (6% of the population in the region is Indigenous)^18^ and is consistent with previous work examining the incidence of IPV in Indigenous populations in Canada.^19–21^

**Table 2.**
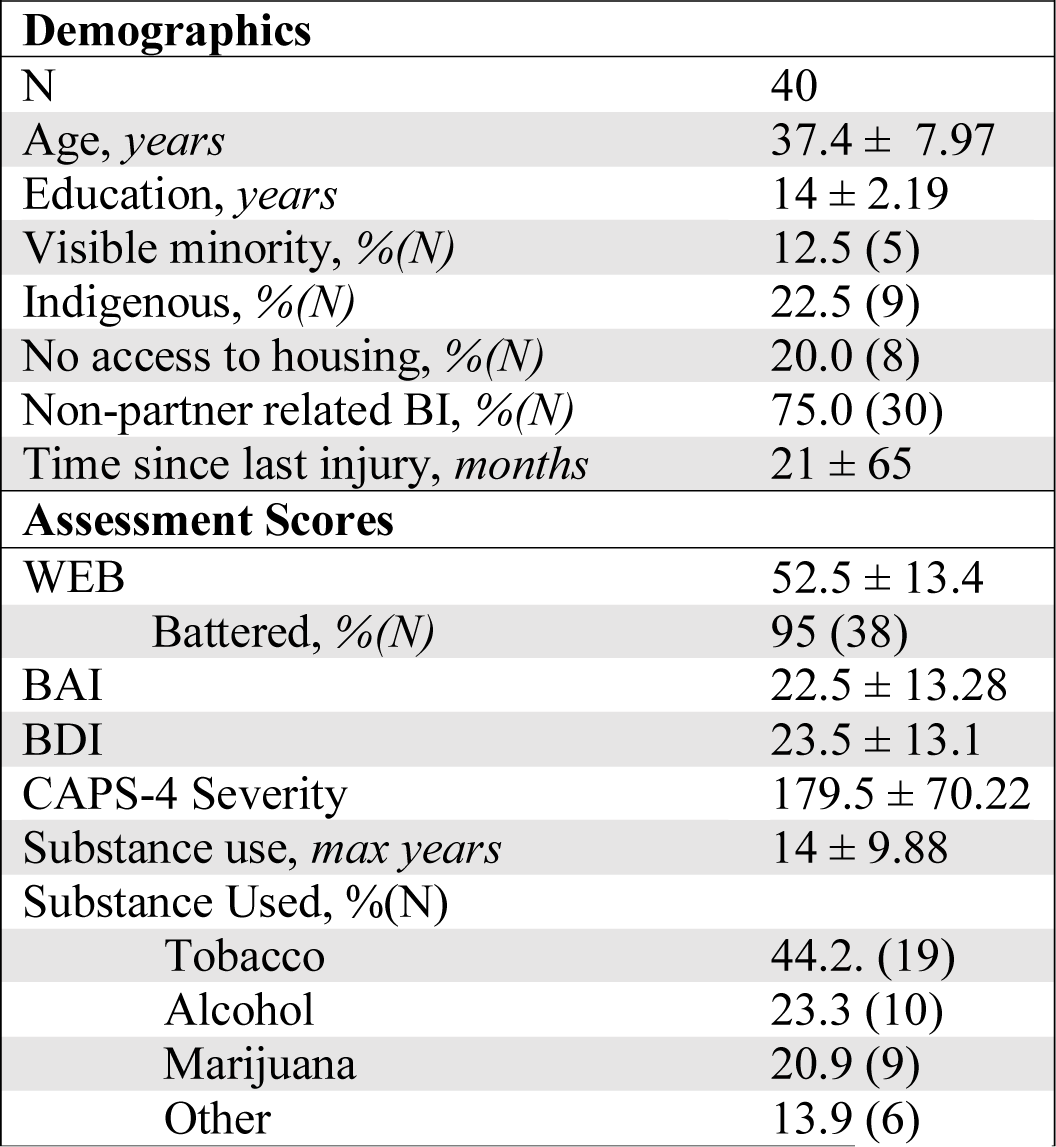
Participant characteristics; median ± standard deviation (SD) unless otherwise specified. WEB: Women’s Experience with Battering scale; BAI: Beck Anxiety Inventory; BDI: Beck Depression Inventory; CAPS-4: Clinician-Administered PTSD Scale for DSM-4

| **Demographics** |  |
| --- | --- |
| N | 40 |
| Age, *years* | 37.4 ± 7.97 |
| Education, *years* | 14 ± 2.19 |
| Visible minority, %(N) | 12.5 (5) |
| Indigenous, %(N) | 22.5 (9) |
| No access to housing, %(N) | 20.0 (8) |
| Non-partner related BI, %(N) | 75.0 (30) |
| Time since last injury, *months* | 21 ± 65 |
| **Assessment Scores** |  |
| WEB | 52.5 ± 13.4 |
| Battered, %(N) | 95 (38) |
| BAI | 22.5 ± 13.28 |
| BDI | 23.5 ± 13.1 |
| CAPS-4 Severity | 179.5 ± 70.22 |
| Substance use, *max years* | 14 ± 9.88 |
| Substance Used, %(N) |  |
| Tobacco | 44.2. (19) |
| Alcohol | 23.3 (10) |
| Marijuana | 20.9 (9) |
| Other | 13.9 (6) |

### Stepwise Multiple Linear Regression

The outputs of the stepwise multiple regression analysis are shown in Table 3. Model fits as assessed with the AIC indicated total targets hit, total objects hit, and right and left hand speed were predicted by the BISA score along with different combinations of PTSD, anxiety, and depression. Moreover, total distractors hit, distractor proportion, and object processing rate were significantly associated with PTSD and depression, but not with the BISA score. Age, history of non-partner related BI, abuse history, and substance use did not significantly contribute to the model and the remaining OHA performance variables (Table 1) were not significantly accounted for by any of the predictors.

**Table 3.**
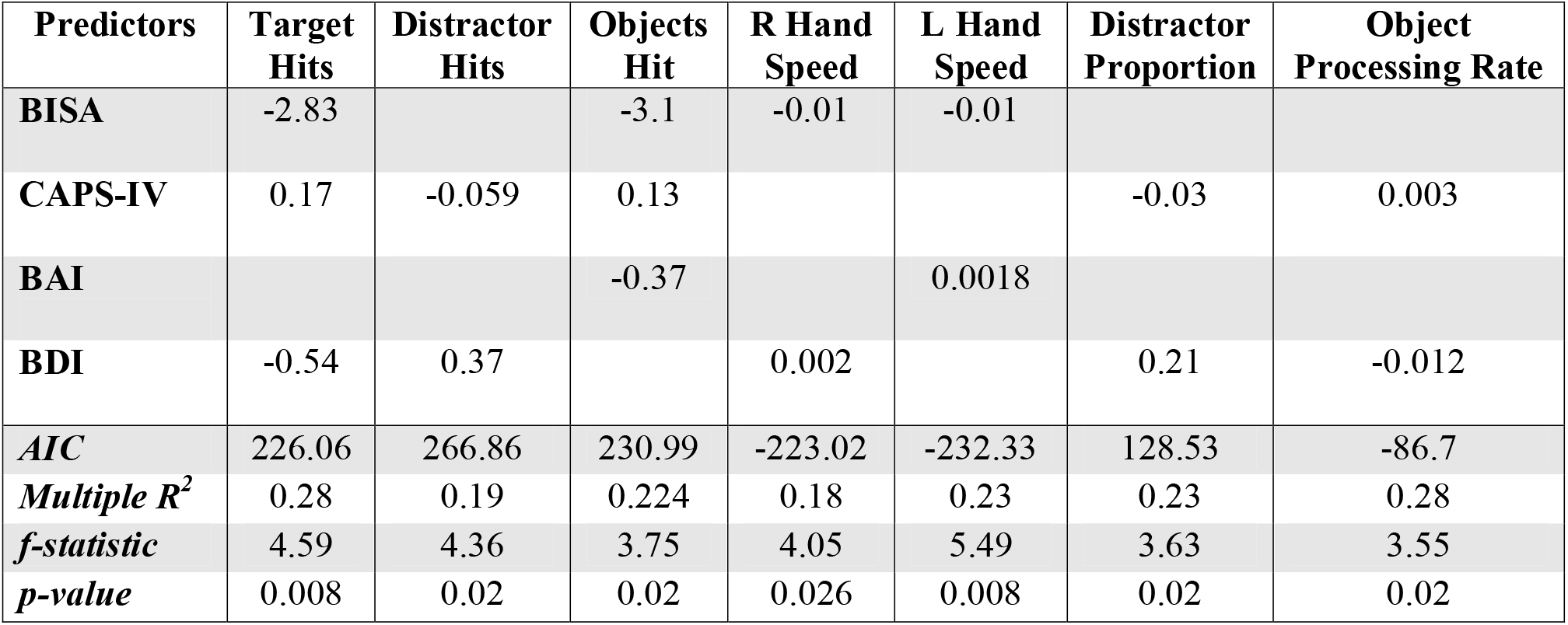
Results of stepwise regression analysis. Each cell represents the significant model estimates for each predictor for each dependent variable. Assumptions of normality, linearity, and homoscedasticity were all met and there were no influential cases. BISA: Brain Injury Severity Assessment; CAPS-IV: Clinician-Administered PTSD Scale; BAI: Beck Anxiety Inventory; BDI: Beck Depression Inventory; AIC: Akaike Information Criterion.

To visualize these results, we plotted simple linear regressions for a subset of the BISA results. Figure 1A shows women with higher BISA scores tended to hit fewer targets compared to those who had lower BISA scores. Post-hoc analysis showed this relationship was driven by the frequency (Figure 1B) and severity (Figure 1C) subcomponents of the BISA, but not the recency. More specifically, women who had experienced 11 or more BIs hit fewer targets than women who experienced 10 or fewer BIs (t-test=1.78, p=0.04). Similarly, women who reported loss of consciousness and/or a period of post-traumatic amnesia following an episode of IPV hit fewer targets than women who did not (t-test=1.89, p=0.03). Additionally, women with higher BISA scores tended to move more slowly relative to those with lower BISA scores (Figure 2A). Post-hoc analysis revealed that this was driven by the recency subcomponent of the BISA (Figure 2B): those who experienced their most recent IPV-related BI more than a year ago had faster hand speeds than women who had experienced an IPV-related BI in the past year (t-test=2.11, p=0.02).

**Figure 1:**
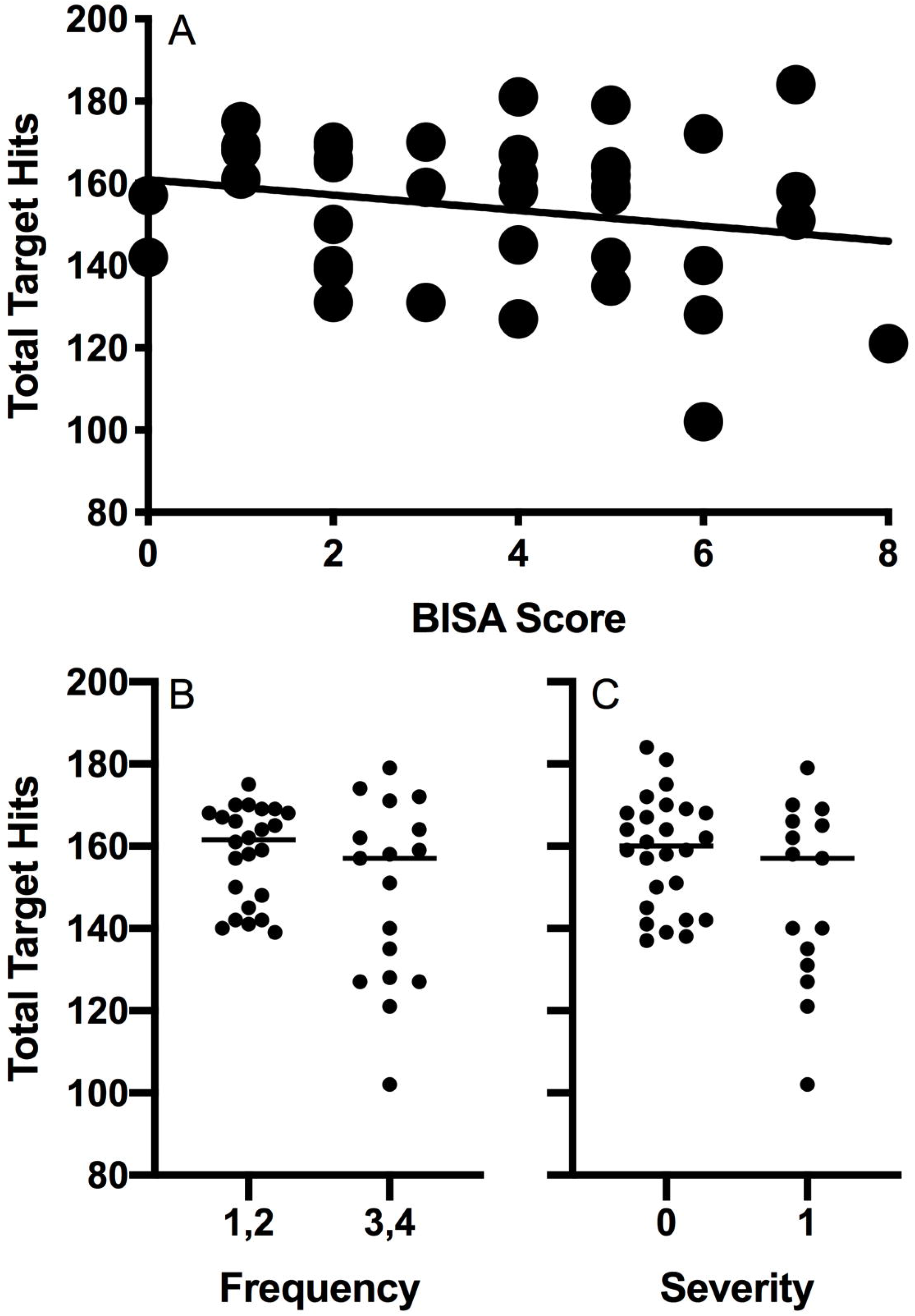
Total target hits plotted as a function of BISA score (A). Total target hits for participants who had experienced 10 or fewer BIs (*frequency scores*: 1,2) vs. those who had experienced 11 or more BIs (*frequency scores*: 3,4) (B). Total target hits for those who had not lost consciousness and/or had a period of post-traumatic amnesia (*severity score*: 0) vs. those who had (*severity score*: 1) (C).

**Figure 2:**
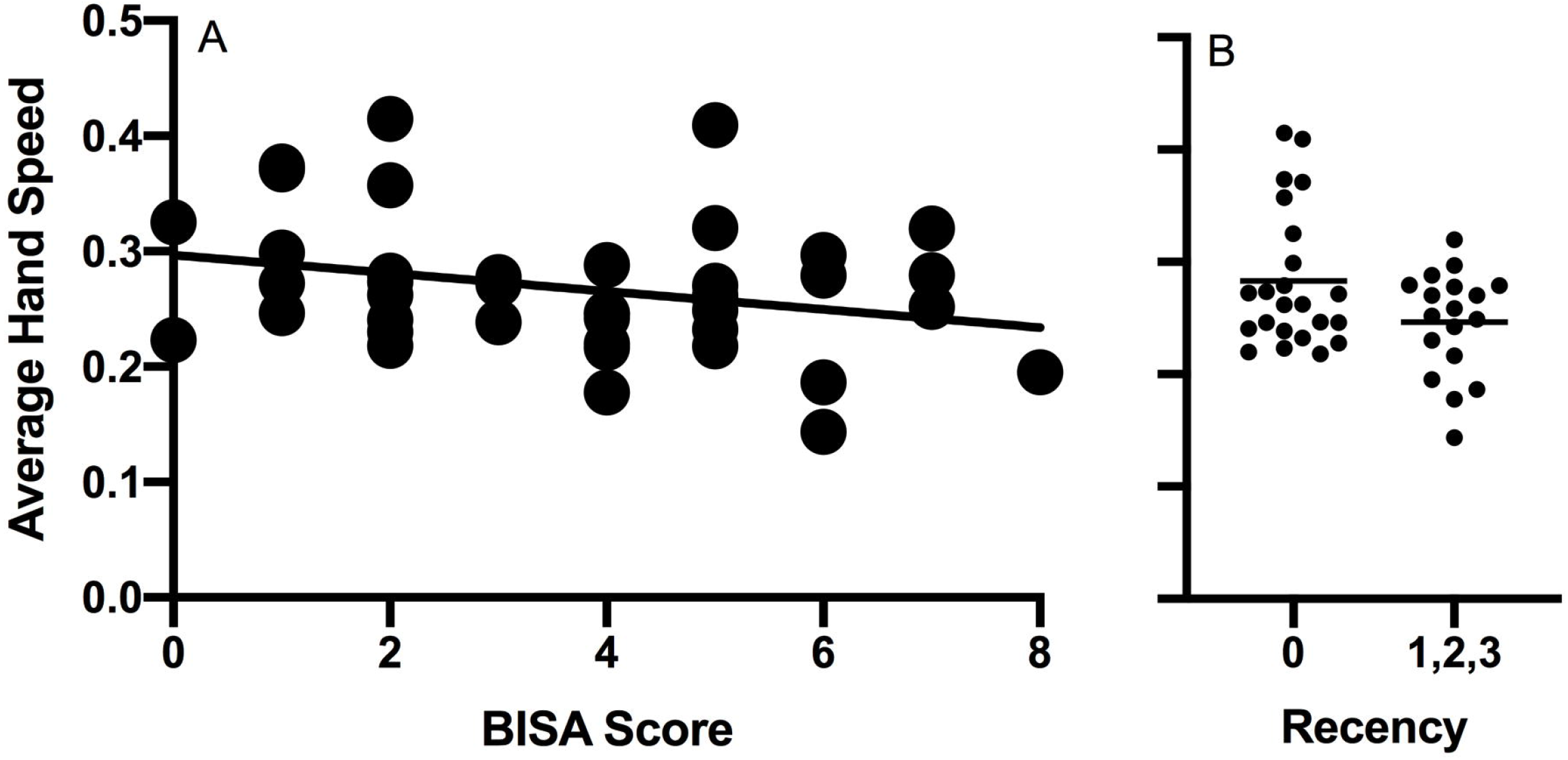
Hand speed (averaged across left and right hands) plotted as a function of BISA score (A). Average hand speed for participants who had experienced their most recent episode more than a year ago (*recency score*: 0) vs. those who had experienced their most recent episode within the last year (*recency scores*: 1,2,3) (B).

### Executive Function Measures

We found significant negative relationships between total target hits and total object hits and trail-making A time (*r* = −0.39, *p* = 0.012; *r* = −0.37, *p* = 0.017) and trail-making B time (*r* = −0.42, *p* = 0.008 for both). No significant correlations were found between these performance measures and BRIEF-A scores or for the hand speed measures.

## DISCUSSION

This exploratory study provides a first investigation of cognitive-motor function in women who have experienced IPV-related BI. Our findings demonstrate IPV-related BI disrupts cognitive-motor function years in that BI load, as assessed by the BISA score, accounted for a significant amount of variability in the number of targets hit, the number of objects hit, and hand speed. PTSD, anxiety, and depression also contributed significantly to measures. In addition, the number and proportion of distractors hit and the object processing rate was significantly driven by PTSD and depression. The fact that comorbidities such as PTSD, anxiety, and depression also contribute to these challenges reflects the complex interaction between the physical injuries induced by the episodes of IPV and the resulting impacts these experiences have on mental health.

Our findings add to the small but growing body of literature demonstrating the chronic and deleterious effects of BI in this population. In particular, the results of this study show women who have experienced IPV exhibit cognitive-motor deficits that can be attributed to BI load as assessed with the BISA. The fact that these deficits were correlated with trail-making task performance suggests that they also offer clinically meaningful information. Moreover, the current results are consistent with studies of repetitive head impacts in collision sport athletes showing impairments in cognitive-motor function that have been linked to an increased risk for long-term impairment.^22^ This is relevant in this context given that survivors of IPV may experience violent episodes at regular intervals over a period of months or years.^23,24^

It remains unknown whether chronic and repetitive exposure to IPV, and, by extension, potential BI, puts women at risk for long-term cognitive impairment or neurodegeneration. It has been demonstrated spousal abuse increases the odds of an Alzheimer’s diagnosis in women^25^ and there is at least one case report of a woman who had experienced IPV with progressive dementia and morphological brain changes consistent with ‘dementia pugilistica’, the historical precursor to chronic traumatic encephelopathy (CTE).^26^ Progressive neurocognitve or sensorimotor deficits in individuals with a history of head impacts have been shown to be a clinical milestone in CTE, and more severe sensorimotor dysfunction is often considered a sign of more advanced neurodegeneration.^27–31^ Crucially, the median time since the last IPV-related BI in our sample was just under 2 years with a substantial range of times across participants. Thus, the deficits observed were very much chronic in nature. In light of the growing body of evidence suggesting high exposure to both concussive and subconcussive impacts may increase the risk of progressive neurodegeneration^32^, there is an urgent need to investigate these hypothesized relationships in the context of IPV.

Several parameters of OHA performance, namely those related to executive dysfunction (number and proportion of distractor hits and the object processing rate), were associated with PTSD and depression. Evidence supporting a relationship between psychopathologies, particularly those triggered by traumatic events, and executive dysfunction is equivocal. Our results may reflect changes in impulsivity as well as reduced inhibitory control – an interpretation consistent with previous studies reporting neurocognitive deficits in both IPV and associated psychopathologies.^8,33,34^ Ultimately, the fact that comorbidities such as PTSD, anxiety, and depression also contribute to these challenges reflects the complex interaction between the trauma – whether emotional, psychological, or physical - induced by episodes of IPV and the resulting impacts these experiences have on cognitive function. From a pathophysiological perspective, it is possible all these factors contribute synergistically to creating a maladaptive physiological environment conducive to cognitive-motor dysfunction. In light of these complexities, it is imperative that resources created to support women who have experienced IPV take a holistic and multidisciplinary approach and that future research efforts examine these interactions to maximize the ecological validity of their findings.

Broadly speaking, the deficits observed in the current lab-based assessment are consistent with reduced executive function and may contribute to the cognitive symptoms reported by women experience IPV^11^. Such deficits may present challenges for IPV survivors in making appropriate and timely decisions around safety planning and accessing supports and, thereby, contribute to difficulties in successfully transitioning to a life free from abuse. Therefore, supports, resources, and training created for this population should consider the added burden of BIs on women experiencing IPV. Whereas anti-racist and anti-discriminatory trauma- and violence-informed care should be at the heart of all health care practices, further knowledge and awareness of IPV-related BI and its effect on brain function may help explain some of the behaviors historically attributed to faults in character.^35^

Despite the novel findings of this research, it should be viewed as a pilot study with a number of limitations. First, due to the small sample size, we may lack sufficient power to detect subtle but meaningful deficits. In addition, the cohort tested was a convenience sample of women recruited from local anti-violence organizations, thus limiting the generalizability of our findings. It is well-documented that IPV can affect all women, irrespective of age, race, ethnicity, and religion; thus, it is essential future studies include a more diverse sample. In addition, by recruited primarily from a shelter that asked residents to remain clean during their stay and asking participants to refrain from consuming alcohol or using drugs for 24 hours prior to testing, we may have excluded women with more severe episodes of IPV and, thereby, failed to encompass a wide spectrum of experiences. Furthermore, the overrepresentation of Indigenous women amongst our participants is consistent with the literature^36^ and highlights the urgent need for more collaborative engagement, education, services, and research in these communities. Haag and colleagues^37^ recently described IPV knowledge and service gaps in several Inuit and First Nations communities – further supporting a call to action and emphasizing the need for a strength-based and collaborative approach in studying IPV with Indigenous communities. It remains critical to study and quantify IPV-related BI in these populations to better understand the systemic barriers to care and safety they face. Given the cross-sectional design of this study, one should also be cautious about drawing causal links between IPV-related BI and cognitive-motor function. Other experiences, such as childhood abuse may also partly account for the alterations in cognitive-motor function we observed. Unfortunately, although participants sometimes referred to such experiences in our assessments, we did not systematically characterize them. Lastly, we did not include a control group, thus limiting our ability to infer the independent effect of IPV-related BI on cognitive-motor function. Indeed, identifying an appropriate control group for IPV-related BI survivors presents a significant methodological challenge. Whereas the ideal control group would consist of women who have experienced IPV but not a BI, our data indicate this is an uncommon occurrence, as evidenced by the fact just 2 of the 40 women (5%) included in the current investigation reported a BISA score of 0 (the score expected from someone who has experienced IPV but not sustained an IPV-related BI). This further highlights the prevalence of IPV-related BIs and underscores the urgency for continued research efforts.

Despite these limitations, this study provides a first exploration of cognitive-motor function related to IPV-related BI and adds to the growing body of literature demonstrating evidence of chronic effects of BI in this population. Future studies should aim to further characterize IPV-related BI in a larger, more heterogeneous sample. In addition, given the association between cognitive impairment and history of BI, there is a need for longitudinal studies documenting neurocognitive and sensorimotor function in women who have experienced IPV to better understand the potential long-term consequences of IPV-related BIs, including the risk for cognitive impairment and neurodegeneration. Ultimately, investigating the physiological basis of IPV-related BI will improve our understanding of the various dimensions of this experience, and support efforts to improve access to resources designed for this population.

## CONCLUSION

Overall, our findings suggest IPV-related BI disrupts the processing required to quickly and accurately hit targets while avoiding distractors in a complex sensorimotor task. The fact that comorbidities such as PTSD, anxiety, and depression also contribute to these challenges reflects the complex interaction between the physical injuries induced by the episodes of IPV and the resulting impacts these experiences have on mental health. Our findings are consistent with previous research demonstrating the long-lasting effects of IPV-related BI and support the call for more trauma-informed BI supports and resources for this underserved population.

## ACKNOWLEDGEMENTS

The authors thank the clients and staff at Kelowna Women’s Shelter and Central Okanagan Elizabeth Fry Society. NMR was supported by a Canada Graduate Scholarship – Master’s award. This work was supported by grants from the Department of Women and Gender Equality, Canadian Institutes of Health Research, Canadian Foundation for Innovation, and an anonymous donor.

## REFERENCES

1. World Health Organization. (2012). Understanding and adressing violence against women.

2. Campbell, J.C., Anderson, J.C., McFadgion, A., Gill, J., Zink, E., Patch, M., Callwood, G., and Campbell, D. (2018). The Effects of Intimate Partner Violence and Probable Traumatic Brain Injury on Central Nervous System Symptoms. J. Women’s Heal. 27, 761–767.

3. Valera, E., and Kucyi, A. (2017). Brain injury in women experiencing intimate partner-violence: neural mechanistic evidence of an “invisible” trauma. Brain Imaging Behav. 11, 1664–1677.

4. Zieman, G., Bridwell, A., and Cardenas, J.F. (2017). Traumatic Brain Injury in Domestic Violence Victims: A Retrospective Study at the Barrow Neurological Institute. J. Neurotrauma 34, 876–880.

5. Valera, E.M., Cao, A., Pasternak, O., Shenton, M.E., Kubicki, M., Makris, N., and Adra, N. (2019). White Matter Correlates of Mild Traumatic Brain Injuries in Women Subjected to Intimate-Partner Violence: A Preliminary Study. J. Neurotrauma 36, 661–668.

6. Hunnicutt, G., Lundgren, K., Murray, C., and Olson, L. (2017). The intersection of intimate partner violence and traumatic brain Injury: A call for interdisciplinary research. J. Fam. Violence 32, 471–480.

7. Haag, H.L., Jones, D., Joseph, T., and Colantonio, A. (2019). Battered and brain injured: Traumatic brain injury among women survivors of intimate partner violence-A scoping review. Trauma. Violence Abuse, 1524838019850623.

8. Daugherty, J.C., Marañón-Murcia, M., Hidalgo-Ruzzante, N., Bueso-Izquierdo, N., Jiménez-González, P., Gómez-Medialdea, P., and Pérez-García, M. (2018). Severity of neurocognitive impairment in women who have experienced intimate partner violence in Spain. J. Forens. Psychiatry Psychol. 30, 322–340.

9. Kwako, L.E., Glass, N., Campbell, J., Melvin, K.C., Barr, T., and Gill, J.M. (2011). Traumatic brain injury in intimate partner violence: a critical review of outcomes and mechanisms. Trauma. Violence Abuse 12, 115–126.

10. Smirl, J.D., Jones, K.E., Copeland, P., Khatra, O., Taylor, E.H., and Van Donkelaar, P. (2019). Characterizing symptoms of traumatic brain injury in survivors of intimate partner violence. Brain Inj. 33, 1529–38.

11. Valera, E.M., and Berenbaum, H. (2003). Brain injury in battered women. J. Consult. Clin. Psychol. 71, 797–804.

12. Belanger, H.G., Spiegel, E., and Vanderploeg, R.D. (2010). Neuropsychological performance following a history of multiple self-reported concussions: A meta-analysis. J. Int. Neuropsychol. Soc. 16, 262–267.

13. Karr, J.E., Areshenkoff, C.N., and Garcia-Barrera, M.A. (2014). The neuropsychological outcomes of concussion: A systematic review of meta-analyses on the cognitive sequelae of mild traumatic brain injury. Neuropsychology 28, 321–336.

14. Karr, J.E., Areshenkoff, C.N., Duggan, E.C., and Garcia-Barrera, M.A. (2014). Blast-Related Mild Traumatic Brain Injury: A Bayesian Random-Effects Meta-Analysis on the Cognitive Outcomes of Concussion among Military Personnel. Neuropsychol. Rev. 24, 428–444.

15. Moore, R.D., Lepine, J., and Ellemberg, D. (2017). The independent influence of concussive and sub-concussive impacts on soccer players’ neurophysiological and neuropsychological function. Int. J. Psychophysiol. 112, 22–30.

16. Mainwaring, L., Pennock, K.M.F., Mylabathula, S., and Alavie, B.Z. (2018). Subconcussive head impacts in sport: A systematic review of the evidence. Int. J. Psychophysiol. 132, 39–54.

17. Breedlove, E.L., Robinson, M., Talavage, T.M., Morigaki, K.E., Yoruk, U., O’Keefe, K., King, J., Leverenz, L.J., Gilger, J.W., and Nauman, E.A. (2012). Biomechanical correlates of symptomatic and asymptomatic neurophysiological impairment in high school football. J. Biomech. 45, 1265–1272.

18. Statistics Canada. (2017). Focus on Geography Series, 2016 Census. Statistics Canada Catalogue no. 98-404-X2016001. Ottawa, Ontario.

19. Brownridge, D.A. (2008). Understanding the Elevated Risk of Partner Violence Against Aboriginal Women: A Comparison of Two Nationally Representative Surveys of Canada. J. Fam. Violence 23, 353–367.

20. Daoud, N., Smylie, J., Urquia, M., Allan, B., and O’Campo, P. (2013). The contribution of socio-economic position to the excesses of violence and intimate partner violence among Aboriginal versus non-Aboriginal women in Canada. Can. J. Public Heal. 104.

21. Pedersen, J.S., Malcoe, L.H., and Pulkingham, J. (2013). Explaining Aboriginal/Non-Aboriginal Inequalities in Postseparation Violence Against Canadian Women: Application of a Structural Violence Approach. Violence Against Women 19, 1034–1058.

22. Montenigro, P.H., Alosco, M.L., Martin, B.M., Daneshvar, D.H., Mez, J., Chaisson, C.E., Nowinski, C.J., Au, R., McKee, A.C., Cantu, R.C., McClean, M.D., Stern, R.A., and Tripodis, Y. (2017). Cumulative Head Impact Exposure Predicts Later-Life Depression, Apathy, Executive Dysfunction, and Cognitive Impairment in Former High School and College Football Players. J. Neurotrauma 34, 328–340.

23. Thompson, R.S., Bonomi, A.E., Anderson, M., Reid, R.J., Dimer, J.A., Carrell, D., and Rivara, F.P. (2006). Intimate Partner Violence Prevalence, Types, and Chronicity in Adult Women. Am. J. Prev. Med. 30, 447–457.

24. Jackson, H., Nuttall, R.L., Philp, E., and Diller, L. (2002). Traumatic brain injury: A hidden consequence for battered women. Prof. Psychol. Res. Pract. 33, 39–45.

25. Leung, F.-H., Thompson, K., and Weaver, D.F. (2006). Evaluating spousal abuse as a potential risk factor for Alzheimer’s disease: rationale, needs and challenges. Neuroepidemiology 27, 13–16.

26. Roberts, G.W., Whitwell, H.L., Acland, P.R., and Bruton, C.J. (1990). Dementia in a punch-drunk wife. Lancet 335, 9180919.

27. McKee, A.C., Stein, T.D., Nowinski, C.J., Stern, R.A., Daneshvar, D.H., Alvarez, V.E., Lee, H.-S., Hall, G., Wojtowicz, S.M., Baugh, C.M., Riley, D.O., Kubilus, C.A., Cormier, K.A., Jacobs, M.A., Martin, B.R., Abraham, C.R., Ikezu, T., Reichard, R.R., Wolozin, B.L., Budson, A.E., Goldstein, L.E., Kowall, N.W., and Cantu, R.C. (2020). The spectrum of disease in chronic traumatic encephalopathy. Brain 136, 43–64.

28. Kornguth, S., Rutledge, N., Perlaza, G., Bray, J., and Hardin, A. (2017). A proposed mechanism for development of CTE following concussive events: Head impact, water hammer injury, neurofilament release, and autoimmune processes. Brain Sci. 7, 1–15.

29. Golden, C.J., and Zusman, M.R. (2019). Chronic Traumatic Encephalopathy (CTE) Impact on Brains, Emotions, and Cognition. Springer.

30. Bhowmick, S., D‘Mello, V., Ponery, N., and Abdul-Muneer, P.M. (2018). Neurodegeneration and sensorimotor deficits in the mouse model of traumatic brain injury. Brain Sci. 8, 12–16.

31. Mckee, A.C., Cantu, R.C., Nowinski, C.J., Hedley-Whyte, T.E., Gavett, B.E., Budson, A.E., Santini, V.E., Lee, H.-S., Kubilus, C.A., and Stern, R.A. (2009). Chronic Traumatic Encephalopathy in Athletes: Progressive Tauopathy After Repetitive Head Injury. J. Neuropathol. Exp. Neurol. 68, 709–735.

32. Mckee, A.C., Alosco, M., and Huber, B.R. (2016). Repetitive Head Impacts and Chronic Traumatic Encephalopathy. Neurosurg. Clin. 27, 529–535.

33. Stricker, N.H., Keller, J.E., Castillo, D.T., and Haaland, K.Y. (2015). The Neurocognitive Performance of Female Veterans With Posttraumatic Stress Disorder. J. Trauma. Stress 28, 102–109.

34. Twamley, E.W., Allard, C.B., Thorp, S.R., Norman, S.B., Cissell, S.H., Berardi, K.H., Grimes, E.M., and Stein, M.B. (2009). Cognitive impairment and functioning in PTSD related to intimate partner violence. J. Int. Neuropsychol. Soc. 15, 879–887.

35. Overstreet, N.M., and Quinn, D.M. (2013). The Intimate Partner Violence Stigmatization Model and Barriers to Help Seeking. Basic Appl. Soc. Psych. 35, 109–122.

36. Native Women’s Association of Canada. (2019). Fact Sheet: Violence Against Aboriginal Women. 1–5 p.

37. Haag, H.L., Sokoloff, S., MacGregor, N., Broekstra, S., Cullen, N., and Colantonio, A. (2019). Battered and Brain Injured: Assessing Knowledge of Traumatic Brain Injury Among Intimate Partner Violence Service Providers. J. women’s Heal. 28, 990–996.

